# Creatine transporter knockout mice (*Slc6a8*) show increases in serotonin-related proteins and are resilient to learned helplessness

**DOI:** 10.1101/641845

**Authors:** Zuhair I. Abdulla, Jordan L. Pennington, Arnold Gutierrez, Matthew R. Skelton

## Abstract

Approximately 20% of adults in the U.S. will experience an affective disorder during their life. While it is well established that serotonin (5-HT) is a crucial factor in mood, impaired cellular bioenergetics are also implicated. Creatine (Cr), through the Cr/Phospho-Cr (PCr) shuttle, maintains high ATP concentrations in the neuron. This system may be implicated in the etiology of affective disorders, as reduced Cr, PCr, and ATP are often seen in the brains of affected patients. To address this issue, Cr transporter (Crt) deficient male mice (*Slc6a8*^−/*y*^) and female mice heterozygous for Crt expression (*Slc6a8*^+/−^) were used to evaluate how a Cr deficient system would alter affective-like behaviors. *Slc6a8*^−/y^ and *Slc6a8*^+/−^ mice had more escapes and faster escape latencies in learned helplessness, indicating a potential resilience to behavioral despair. Elevated zero maze and tail-suspension test performance matched that of wildtype mice, however. *Slc6a8*^−/*y*^ mice have increased 5-hydroxyindoleacetic acid content in the hippocampus and striatum and increased monoamine oxidase protein and tryptophan hydroxylase-2 protein content in the hippocampus, while serotonin levels are unchanged. This indicates an increase in 5-HT turnover. Our results indicate that Cr plays a complex role in affective disorders and 5-HT neurotransmission, warranting further investigation.

## INTRODUCTION

Affective disorders, such as depression or anxiety, are a significant worldwide healthcare burden, with depression being the leading cause of disability in the United States (NIMH 2010a; NIMH 2010b). Though primarily thought of as imbalances or deficiencies in monoaminergic activity, alterations to brain bioenergetics have also been implicated in these disorders (Gardner & Boles 2011; Klinedinst & Regenold 2015). Creatine (Cr) is an important spatial and temporal ATP buffer in cells with a high-energy demand (Wyss 2000). Cr is transported into cells via the Cr transporter (Crt; SLC6A8) (Braissant *et al.* 2011; Braissant *et al.* 2001; Hanna-El-Daher & Braissant 2016; Joncquel-Chevalier Curt *et al.* 2015; Perna *et al.* 2016; Skelton *et al.* 2011). Supporting the role of Cr in bioenergetic pathways, Crt knockout (*Slc6a8*^−/*y*^) mice, which lack brain Cr have reduced brain ATP levels (Perna *et al.* 2016; Stockebrand *et al.* 2016). The role of Cr on affective-type behaviors is not fully understood, though changes in brain Cr have been implicated in mood disorders (Allen 2012). The *Slc6a8*^−/*y*^ mice provide a unique opportunity to better understand the role of Cr in affective disorders, by examining the effects of a complete deficit of Cr.

Most studies related to Cr and affective behaviors have been performed following Cr supplementation. In mice, acute Cr administration reduced immobility in the tail-suspension test (TST) (Cunha *et al.* 2016; Cunha *et al.* 2014; Cunha *et al.* 2012; Cunha *et al.* 2015; Cunha *et al.* 2013a; Cunha *et al.* 2013b; Pazini *et al.* 2016). Additionally, Cr supplemented female rats had an increased latency to immobility in the forced-swim test, while Cr-treated males were unaffected (Allen *et al.* 2012). Psychosocial defeat reduced brain Cr in tree shrews (Czeh *et al.* 2001; van der Hart *et al.* 2002). In humans, Cr supplementation improved Hamilton Depression Rating Scale (HAM-D) scores in individuals with treatment-resistant depression (Roitman *et al.* 2007). Combination treatment with escitalopram and Cr increased HAM-D scores and led to a more rapid onset of symptom relief than women treated with a selective serotonin reuptake inhibitor (SSRI) alone (Lyoo *et al.* 2012). Cr levels are reduced in the cerebral white matter of individuals with generalized anxiety disorder and an early life trauma (Coplan *et al.* 2006). Similar findings were reported in the hippocampi of patients with post-traumatic stress disorder (Villarreal *et al.* 2002; Schuff *et al.* 2008). Although a link between diminished levels of Cr and affective disorders is well established, it has yet to be shown what effect a lack of Cr will have on affective behaviors.

The purpose of this study was to determine if changes in affective-like behaviors occur in male *Slc6a8*^−/*y*^ mice and female *Slc6a8*^+/−^ mice. The female mice have heterozygous *Slc6a8* expression and 50% reductions in brain Cr and mild cognitive deficits (Hautman *et al.* 2014). Thusly, we evaluated the effects of a complete or partial lack of Cr in the brain on behaviors related to anxiety and depression. In addition, we evaluated the effect of a loss of Cr on the expression of proteins related to 5-HT synthesis and turnover as well as the locomotor response to a 5-HT based stimulant, para-chloroamphetamine.

## METHODS

### Subjects

Male *Slc6a8*^−/*y*^ and female *Slc6a8*^+/−^ mice, along with wild-type littermate controls were generated in-house by breeding male *Slc6a8*^+/*y*^ with female *Slc6a8*^+/−^ mice. These mice have been maintained on their parent C57BL6/J strain with SNP analysis showing 97% similarity to the stock C57BL6/J line at Jackson Laboratory. Homozygous knockout female mice cannot be generated since *Slc6a8*^−/*y*^ mice do not breed. Mice were genotyped by PCR as previously described (Perna *et al.* 2016; Skelton *et al.* 2011). Mice were housed at 21° C in microisolators 2-4/cage with *ad libitum* access to food and water. Lights were maintained on a 14 h:10 h light:dark cycle with lights on at 0700. All procedures on mice were performed at the Cincinnati Children’s Research Foundation vivarium which is fully accredited by AAALAC International and protocols were approved by the Institutional Animal Care and Use Committee, protocol #2017-0019. All institutional and national guidelines for the care and use of laboratory animals were followed. No randomization was performed to allocate subjects in the study. Adult mice aged 60 to 80 days were assigned by litter to one of three behavioral cohorts, with only one animal per genotype per litter used. The first cohort was tested in elevated zero maze (EZM) and tail-suspension test (TST) in that order. Cohort 2 were assessed for learned helplessness (LH), while a third cohort was used for the PCA challenge. Only tissue from cohort 1 was used for protein analyses. Data were collected from a total of 199 mice, with no sample calculations having been performed; instead sample sizes were determined by previously collected data. A flow chart showing how animals were used is included as supplemental information. No exclusion criteria were predefined. All behavioral testing was run between 1000 and 1400 on mice.

### Elevated Zero Maze

The protocol followed those previously published with minor modifications (Amos-Kroohs *et al.* 2013; Schaefer *et al.* 2009; Sprowles *et al.* 2016). The maze consists of a circular runway [61 cm (dia) × 5.1 cm (width), 48.5 cm above the ground] with two walled corridors (inner wall [16 cm (h) × 42 cm (l)] and outer walls [16 cm (h) × 49 cm (l)]) on opposite sides, creating alternating open and closed quadrants. The room lights were dimmed to 26.5 lux. Mice were placed in a closed quadrant and the experimenter left the room. Using a closed-circuit digital camera system, behavior was scored in the hallway outside of the procedure room. Latency to the first open quadrant entrance, total time spent in open quadrants, number of distinct open quadrant entries, and number of head dips were live scored by an experimenter blinded to the treatment using ODLog software (Macropod Software).

### Tail-Suspension Test

The tail-suspension apparatus consists of a transparent platform [33 cm (l) × 20 cm (d) x 16.5 cm (h)] with 4 holes (0.635 cm diameter) covered by rubber grommets (Schaefer *et al.* 2009). At the beginning of the 5 min (300 s) trial, the mouse’s tail was passed through a hole from the underside of the platform and the mouse was raised to the point where it was unable to reach up and grasp the base of its tail for support. The tail was taped to the top of the platform. Trials were live-scored for latency to first immobility, total mobility time, and total immobility time (Can *et al.* 2012).

### PCA Challenge

Baseline and stimulant-induced locomotor activity was monitored using the PAS system from San Diego Instruments (San Diego, CA). The 41 × 41-cm chambers are made of clear acrylic and are lined with horizontal strips of 16 LED infrared beams, with each beam spaced 2.56 cm apart. Mice were placed in the chambers for 30 min to measure baseline activity. Following baseline, mice received an i.p. injection of saline (10 ml/kg) and were placed back into the chamber for 30 min to control for injection stress. Mice then received 1 mg/kg, 2.5 mg/kg, or 5 mg/kg of the 5-HT agonist *para*-chloroamphetamine (PCA; volume 10 mg/ml, adjusted for free base) and were placed back into the arena for 2 h. Unique and repetitive beam breaks were recorded by the PAS software. Total ambulatory activity is calculated by taking the sum of all unique beam breaks. Data are analyzed as the sum of total activity within 5 min blocks.

### Learned Helplessness

LH was tested using the GEMINI Avoidance System (San Diego Instruments, San Diego, CA). The apparatus consists of an arena with two chambers [size of each chamber: 24 cm (w) × 20.5 cm (d) × 20.5 cm (h)] separated by a mechanical gate. To ensure that the paradigm induced LH, mice were assigned to one of two training protocols: Shock or No-shock. On the two training days, mice were placed in the chamber with the gate closed for approximately 45 min. Mice in the Shock group received 360 foot-shocks (0.3 mA, 2 s duration, randomized intertrial intervals (ITI) of 1–10 s) that were omitted in the No-Shock group. On day three, both groups of mice were tested for learned helplessness. Mice were placed in the original chamber to start the trial; the gate was lifted and a continuous foot shock (0.3 mA) was delivered until the mouse escaped to the adjacent chamber or until the time limit (24 s) had elapsed. Mice received 30 trials with a 10 s ITI. Latency to escape and trials with a successful escape were recorded.

### High Performance Liquid Chromatography

Mice tested in the EZM and TST were used for analysis of neurotransmitter content. Within 1 week of testing, mice were anesthetized with isoflurane, decapitated, and the brain was rapidly removed. The same experimenter performed all dissections to control for the time from dissection to freezing. The hippocampus and striatum tissue were dissected free-hand, flash frozen in dry-ice cooled methylbutane, and stored at −80° C until processing. Tissue was sonicated in ice-cold 0.1 N perchloric acid (300 µl per striatum, 500 µl per hippocampus) and centrifuged at 20,800 RCF for 13 min at 4.0° C. Monoamine concentrations were measured from the supernatant using high-performance liquid chromatography (HPLC). The HPLC-ECD system consists of a Dionex UltiMate^®^ 3000 Analytical Autosampler connected to an ESA 5840 pump and a Coulochem III electrochemical detector (Thermo Scientific), which was set to −150 mV for E1 and +250 mV for E2. The guard cell was set to +350 mV. The column used was a Supelco Supelcosil™ LC-18 column (15 cm × 4.6 mm, 3 μm; Sigma-Aldrich Co.). MD-TM Mobile Phase (Thermo Fisher Scientific; 89% water, 10% acetonitrile, and 1% sodium phosphate monobasic (monohydrate)) was used. The flow rate was 0.5 ml/min. Serial dilutions of standards from each monoamine were used to generate a standard curve and determine the concentration of the tissue samples.

### Western Blots

Hippocampi were homogenized by sonication in RIPA buffer with protease inhibitors (composed of sodium chloride, sodium dodecyl sulfate, sodium deoxycholate, Tris-HCL, Triton X). Protein concentrations were determined using the Pierce bicinchoninic acid assay (BCA; Thermo-Fisher). Samples were mixed with 2x Laemmli buffer (Life Sciences) to a final concentration of 1 μg/μL and incubated at 95° C for 5 min. A total of 10 µg protein/well was separated by electrophoresis on 12% Mini-PROTEAN TGX gels (Bio-Rad). Proteins were transferred to Immobilon-FL PVDF membranes (Millipore) followed by incubation in Odyssey blocking buffer (LI-COR, Lincoln, NE) for 1 h at room temperature (RT). Blots were incubated with antibodies for TPH-2 (Abcam, ab184505), monoamine oxidase A (MAO; Abcam, ab126751), α-tubulin (Sigma Aldrich, T5168), and ß-actin (LI-COR, #926-42212) overnight with agitation at 4° C. Membranes were washed with TBST and incubated at RT with IRDye (LI-COR) secondary antibodies for 1 h. Membranes were again washed with TBS-T followed by TBS and visualized using an Odyssey CLx scanner (LI-COR). Bands were analyzed in Image Studio (LI-COR). TPH-2 and MAO band intensities were normalized to ß-actin.

### Statistics

Male and female mice have distinct genotypes, so using sex as a between-subjects factor would create an imbalanced design as there are no heterozygous males and no homozygous females. Therefore, each sex is analyzed separately with genotype as the only between subjects factor. Statistical analysis was performed using GraphPad Prism (San Diego, CA) software. For experiments with a single variable and where the two groups were compared, unpaired Student’s t-tests were used. For tests involving measurement of multiple time blocks, e.g. locomotor activity and learned helplessness, a repeated-measures analysis of variance (ANOVA) was used with gene as the between subjects factor and time as the repeated factor. Significant effects were analyzed using the False Discovery Rate method of Benjamini, Krieger, and Yekuteli (Benjamini *et al.* 2006). Appropriate nonparametric analyses were used when data did not fit a Gaussian distribution and are noted in the text. The null hypothesis that the groups are the same was rejected at *p*<0.05 for all tests. Outliers were defined as values lying more than three standard deviations from the mean in either direction. Values defined as outliers were removed from analysis.

## RESULTS

### Weights

*Slc6a8*^−/*y*^ mice weighed less than *Slc6a8*^+/*y*^ mice at the start of testing ((MEAN±SEM: *Slc6a8^+/y^* = 24.8±0.3 g; *Slc6a8*^−/*y*^ =15.5±0.4 g); t_36_=19.12. *p*<0.0001). No differences were observed between the female *Slc6a8*^+/+^ (MEAN±SEM=20.1±0.35 g) and *Slc6a8*^+/−^ (MEAN±SEM=19.19±0.35 g) mice.

### Elevated Zero Maze

In males, there was no effect of gene for any variable examined. Female *Slc6a8*^+/−^ mice had more entries into the open arms than *Slc6a8*^+/+^ mice (t_38_=2.153, *p*<0.05; Figure 1c). No differences were observed for other variables. 4 outliers in *Slc6a8*^+/*y*^ and 3 each in *Slc6a8*^−/*y*^, *Slc6a8*^+/+^, and *Slc6a8*^+/−^ were discovered in the latency to open variable and were removed from all EZM analysis.

**Figure 1.**
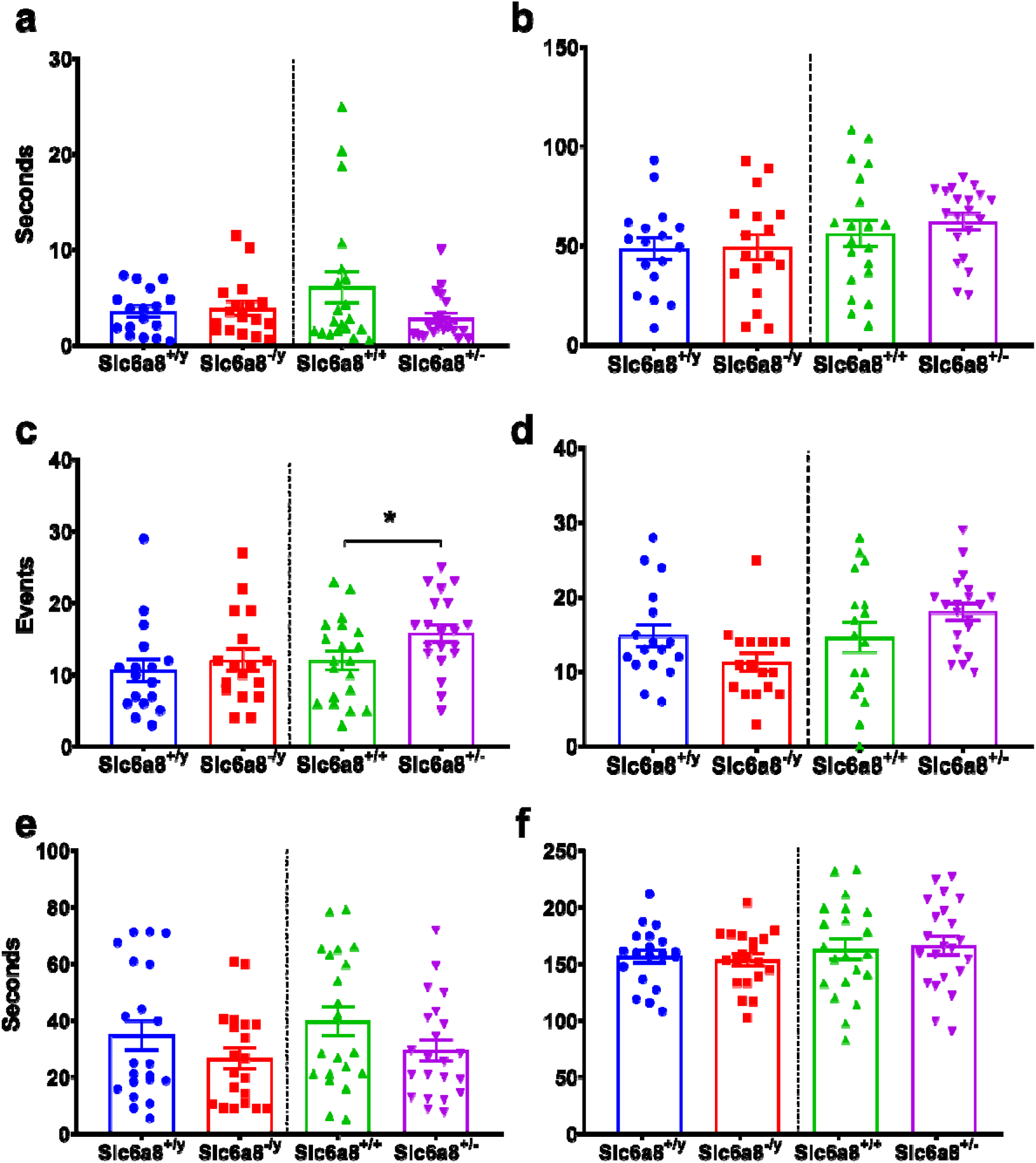
Elevated zero maze and tail-suspension test. (a) Latency to open quadrant entry, (b) time in open quadrants, and (d) head dips were unchanged. (c) *Slc6a8*^+/−^ mice entered the open quadrants more often than *Slc6a8*^+/+^ mice (t_38_=2.153, *p*<0.05). There were 4 outliers in *Slc6a8*^+/*y*^ and 3 each in *Slc6a8*^−/*y*^, *Slc6a8*^+/+^, and *Slc6a8*^+/−^ n=17-20/group. No difference in (e) latency to first immobility or (f) total mobility were observed in *Slc6a8*^−/*y*^ or *Slc6a8*^+/−^ mice. One outlier each was removed from each of the female groups. n=20-22/group, * *p*<0.05.

### Tail-Suspension Test

No effect was observed in either latency to immobility (males: t_39_=0.17, *p*>0.05; females: t_41_=0.53, *p*>0.05; Figure 1d) or mobility time (males: t_39_=0.37, p>0.05; females: t_41_=0.27, *p*>0.05; Figure 1f).

### Learned Helplessness

Mice in the Shock group had longer escape latencies compared with the No Shock group [male (F(1,43)=33.62, *p*<0.0001), female (F(1,51)=12.05, *p*<0.01); Figure 2a], showing that the test paradigm induced learned helplessness. In male mice, there were significant main effects of gene (F(1,22)=6.149, *p*<0.05) and trial (F(29,638)=2.233, *p*<0.001) for latency to escape. *Slc6a8*^−/*y*^ mice also had more trials with an escape *Slc6a8*^+/*y*^ mice (data failed to D’Agostino-Pearson normality test; K2=6.7, *p*<0.05; main effect of gene Mann-Whitney U=28, n_1_=12, n_2_=12, *p*<0.01; Figure 2b). For female mice, there were main effects of gene (F(1,29)=5.37, *p*<0.05) and trial (F(29,841)=3.541, *p*<0.0001) in latency to escape. *Slc6a8*^+/−^ mice had shorter escape latencies than *Slc6a8*^+/+^ mice (Figure 2a). A similar main effect of gene (Mann-Whitney U=62, n_1_=16, n_2_=15, *p*<0.05; Figure 2b) was seen in the number of trial with an escape with the *Slc6a8*^+/−^ mice escaping more than *Slc6a8*^+/+^ mice.

**Figure 2.**
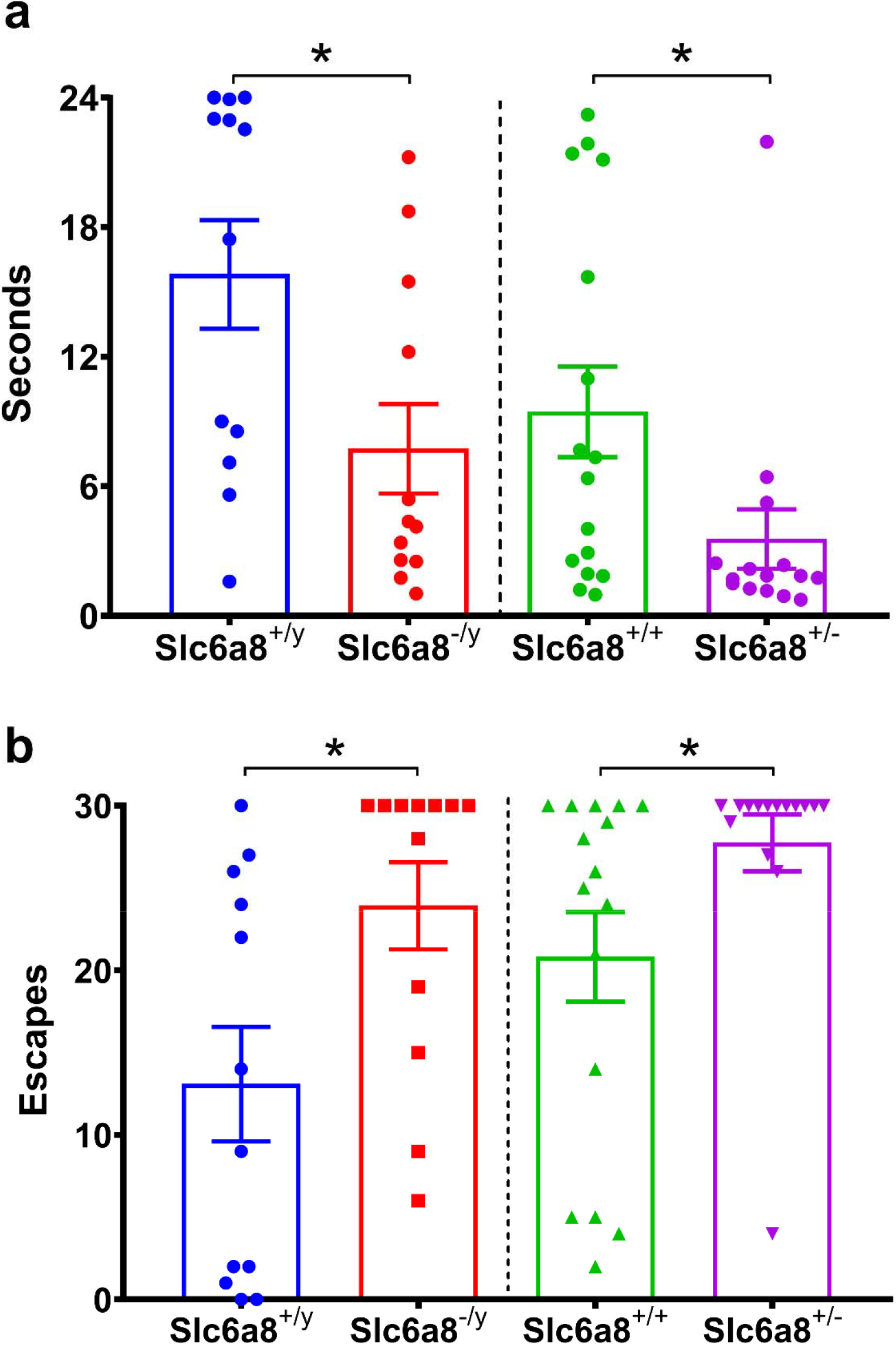
Cr deficient mice are less likely to undergo LH-induction than wild type mice. *Slc6a8*^−/*y*^ and *Slc6a8*^+/−^ mice had decreased latencies to escape (a) and more successful trials (b) than their wild type counterparts. n=12-16/group.

### PCA Challenge

PCA administration at 5 mg/kg produced hyperactivity in both groups (main effect of drug, saline vs PCA; F(1,48)=207.5, p<0.0001; Figure 3). *Slc6a8*^+/*y*^ and *Slc6a8*^−/*y*^ did not differ on any measures. PCA dosed at 1 mg/kg and 2.5 mg/kg failed to produce hyperactivity.

**Figure 3.**
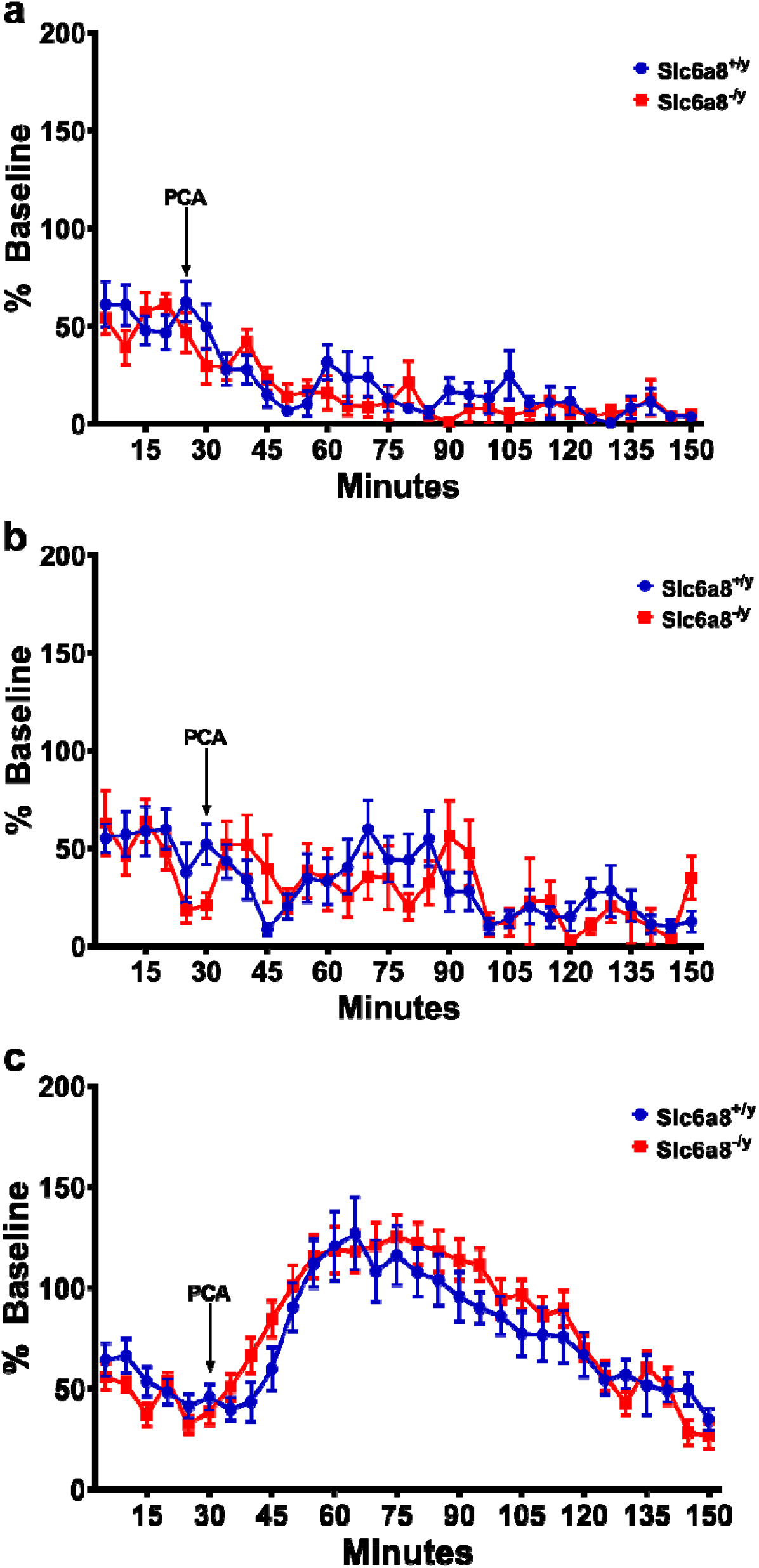
PCA does not uniquely alter the behavior of Cr deficient mice. Neither baseline locomotor activity nor activity in response to PCA were affected. PCA dosed at (a) 1 mg/kg, n=7-8/group, and (b) 2.5 mg/kg, n=8/group, failed to produce hyperactivity, but (c) 5 mg/kg produced robust hyperactivity in both groups, n=13/group.

### High Performance Liquid Chromatography

Males: Hippocampal 5-HIAA levels were increased in *Slc6a8*^−/*y*^ mice compared with *Slc6a8*^+/*y*^ mice (t_22_=2.384, p<0.05; Figure 4b). A similar increase was seen in the neostriatum for 5-HIAA (t_21_=2.488, *p*<0.01; figure 4e). In addition, increases in striatal DA (t_21_=2.493, *p*<0.05; Figure 4g) and 3-HVA (t_21_=2.246, *p*<0.0356; Figure 4i) were observed in *Slc6a8*^−/*y*^ mice compared with *Slc6a8*^+/*y*^ mice. All other neurotransmitters evaluated were unchanged. Females: No differences were found in either brain region for any neurotransmitter or their metabolites.

**Figure 4.**
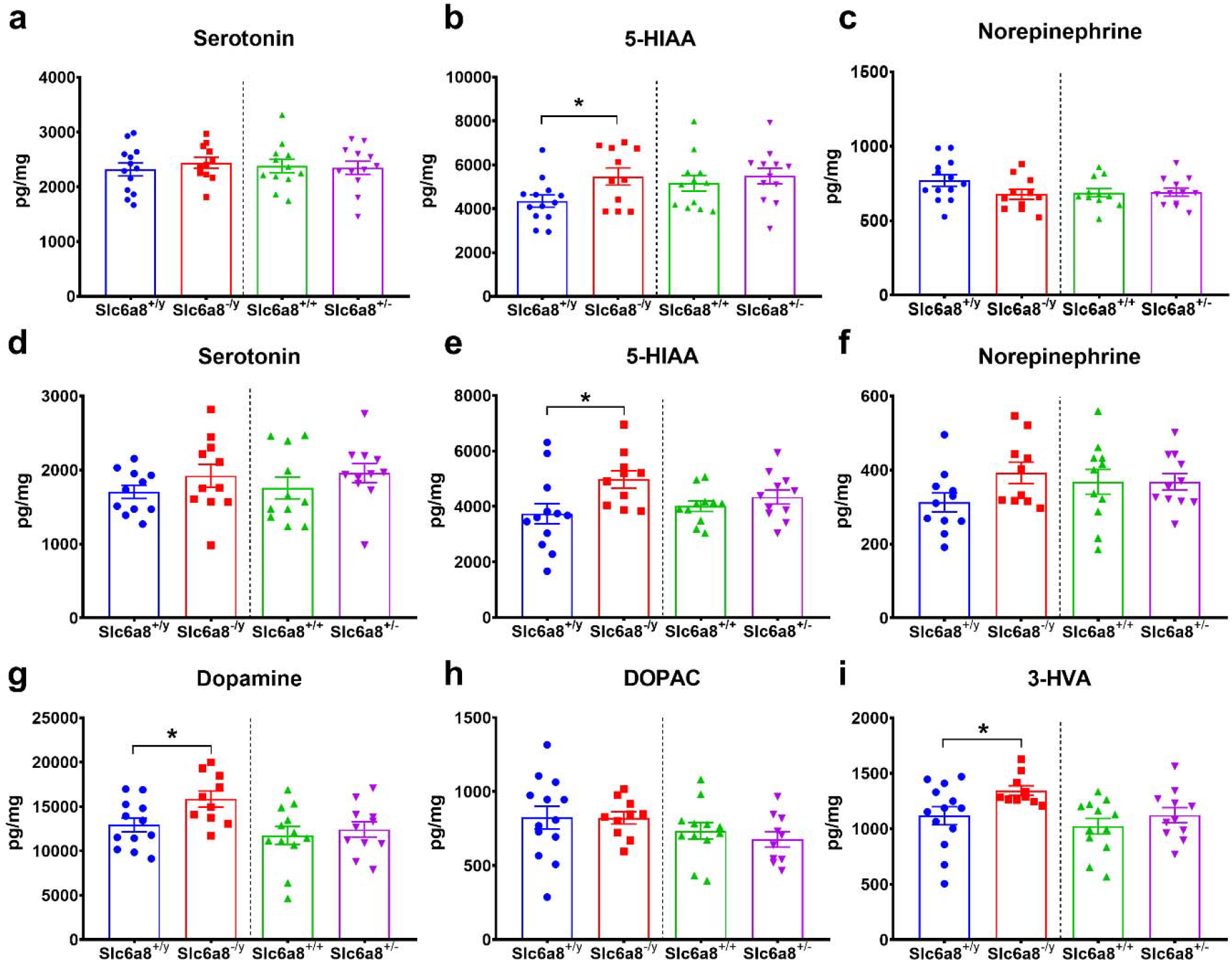
Lack of Crt causes significant alterations to the serotonergic system. (a-c) HIPPOCAMPUS: *Slc6a8*^−/*y*^ mice have increased 5-HIAA concentration (t_22_=2.384, p<0.05; b), but unchanged 5-HT and norepinephrine concentrations. (d-i) STRIATUM: *Slc6a8*^−/*y*^ have increased 5-HIAA (t_22_=2.817, p<0.01, 1 knockout outlier removed; e), dopamine (t_20_=2.493, *p*<0.05, 1 outlier per group removed; g), and increased 3-HVA (t_21_=2.246, *p*<0.05, 1 knockout outlier removed; i) content relative to wild type controls. However, 5-HT concentration is unchanged (d). n=11-13/group.

### Western Blots

Males: TPH-2 signal intensity was increased in the hippocampus of *Slc6a8*^−/*y*^ mice compared with *Slc6a8*^+/*y*^ mice (t_21_=1.872, *p*<0.05, Figure 5a). Hippocampal MAO signal intensity was increased in *Slc6a8*^−/*y*^ vs *Slc6a8*^+/*y*^ mice (t_23_=1.93, *p*<0.05; Figure 5b). Tissue from the striatum was not analyzed. Females: No differences were observed in TPH-2 or MAO expression levels.

**Figure 5.**
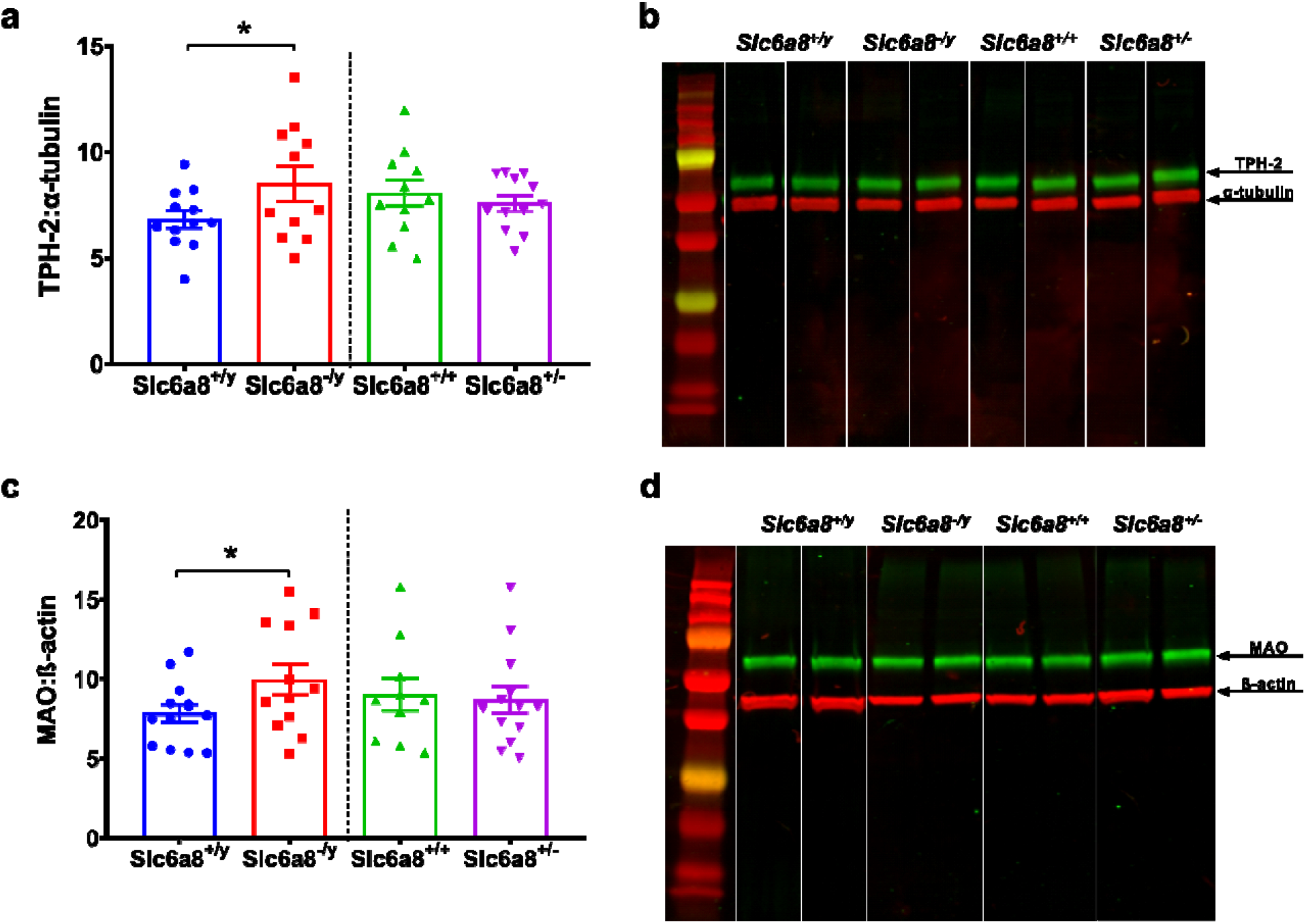
Serotonin-related enzymatic content is increased in Crt-deficient mice. (a) *Slc6a8*^−/*y*^ mice have a greater concentration of TPH-2 in the hippocampus relative to *Slc6a8*^+/*y*^ controls (t_21_=1.872, p<0.05, 1 wildtype outlier removed; (b) Representative composite of TPH-2. (c) *Slc6a8*^−/*y*^ mice also have a greater concentration of MAO in the hippocampus relative to WT controls (t_23_=1.93, p<0.05). (d) Representative composite of MAO westerns. n=10-13/group.

## DISCUSSION

The purpose of this study was to evaluate affective-like behaviors in mice Crt-deficient mice. *Slc6a8*^−/*y*^ and *Slc6a8*^+/−^ mice were resistant to LH, an induced depressive-like behavior. Additionally, male *Slc6a8*^−/*y*^ mice have increased 5-HIAA and increased expression of TPH-2 and MAO, proteins related to 5-HT synthesis and metabolism. Performance in anxiety-like or native depressive-like behavior, as evaluated by the EZM and TST respectively, were not affected by Crt-deficiency. Together, the results of this study suggest that Cr plays a role in 5-HT metabolism, which may manifest as changes in induced depressive-like behaviors.

The LH task represents an animal model of coping strategies that has a high degree of translational value related to depression as it is seen in humans and rodent models (Maier & Seligman 2016). Accordingly, treatment with antidepressants leads to resilience against LH, providing further face validity to this model (Chourbaji *et al.* 2005). The 5-HT system has been shown to play an important role in the acquisition of LH (Maier & Seligman 2016). Additionally, Lugenbiel et al. (2010) found that rats bred for a congenital LH response (cLH) without induction have decreased *Slc6a8* expression in the hippocampus, but not the prefrontal cortex (Lugenbiel *et al.* 2010). Both escitalopram and electroconvulsive shock administration are antidepressant treatments that attenuate LH-responses (Reed *et al.* 2009; Sartorius *et al.* 2003). Interestingly, Lugenbiel at al. demonstrated that these treatments increased *Slc6a8* expression in both the hippocampus and prefrontal cortex of cLH, but not WT, rats. Conversely, this study found that mice with either a lack of or a decreased *Slc6a8* expression were more resistant to LH-induction than wild type mice. While the results of this study do not match the results of the Lugenbiel study, there are caveats that could prevent a direct comparison of these studies. While *Slc6a8* expression was reduced in the hippocampus, Cr levels were not measured in the cLH rats. In addition, the expression of *Slc6a8* was unchanged in the PFC. It is possible that the differences in these studies could be a result of changes caused by a lack of Cr in the PFC. Finally, *Slc6a8* expression was not completely eliminated in the rats tested while it is absent in the *Slc6a8*^−/*y*^ mice. It is possible that reductions in *Slc6a8* and elimination of the gene lead to distinct phenotypes.

Although *Slc6a8*^−/*y*^ and *Slc6a8*^+/−^ mice appear to be more resilient to LH-induction, there were no changes in the tail suspension task or in the zero maze-acute measures of depressive- and anxiety-like behaviors. These seemingly divergent results are not altogether surprising as these tests measure different affective outcomes. A crucial difference between LH and our other affective measurements is that TST and EZM were measured in mice that were not exposed to a known stressor prior to testing. This was done to ensure that any potential alterations in behavior would provide insight into the baseline affective behaviors in mice lacking Slc6a8. In wild-type mice, a depressive-like phenotype is often induced by exposure to known stressors prior to testing (Hammels *et al.* 2015). Based on the differences in the tasks used in this study, it may be beneficial to employ these strategies in *Slc6a8*^−/*y*^ or *Slc6a8*^+/−^ mice in future studies.

We have shown that *Slc6a8*^−/*y*^ and *Slc6a8*^+/−^ mice have cognitive deficits that are caused by the lack of brain Cr (Skelton *et al.* 2011; Udobi *et al.* 2018). It is important to determine what other behaviors may be disrupted by the lack of Cr. In the initial characterization of this model, social interaction and acoustic startle/pre-pulse inhibition were not changed in *Slc6a8*^−/*y*^ mice (Skelton *et al.* 2011). The results from the TST and EZM further establish that the loss of Cr does not lead to a global behavioral deficit. These findings allow for a more precise determination of pathways and brain regions that could be disrupted in the Cr-deficient brain.

It is well-established that LH is dependent on 5-HT function. In the mice used for the initial characterization in Skelton et al, there was a small (11%) increase in hippocampal 5-HT levels of *Slc6a8*^−/*y*^ mice along with a more pronounced (34%) increase in 5-HIAA. In this study, while there was a small increase (5%) in the mean 5-HT content of the hippocampus, it did not reach significance. Given the magnitude of the change between the studies, it would not be surprising that a type-1 error was present in the 2011 study. The increase in 5-HIAA was replicated between the studies with *Slc6a8*^−/*y*^ mice having a 25% increase in hippocampal 5-HIAA levels. The limited or lack of change in 5-HT along with the increase in 5-HIAA would suggest that 5-HT turnover is increased in *Slc6a8*^−/*y*^ mice. Accordingly, we observed an increase in MAO protein levels in the hippocampus of *Slc6a8*^−/*y*^ mice. Additionally, the levels of hippocampal TPH-2 were increased in *Slc6a8*^−/*y*^ mice. The unchanged 5-HT content combined with the elevated TPH-2 in mice lacking Cr suggests that a compensatory mechanism exists in these mice to increase 5-HT levels to accommodate for an increase in 5-HT release and metabolism. Taken together, the increases in 5-HIAA, MAO, and TPH-2 suggests that the loss of *Slc6a8* causes an increase of 5-HT usage and turnover. Although they have a 50% reduction in brain Cr, similar serotonergic alterations were not observed in female *Slc6a8*^+/−^ mice. Therefore, female mice were not included in the PCA challenge.

Acute doses of PCA (0.5 to 5 mg/kg) increase synaptic 5-HT and induce hyperactivity in mice (Fuller 1992; Geyer 1996). Interestingly, there were no significant alterations to motor activity in *Slc6a8*^−/*y*^ mice after PCA administration. It is possible that the high dose of 5 mg/kg PCA caused a ceiling effect with maximal 5-HT release. However, dosing at 1 mg/kg and 2.5 mg/kg failed to produce hyperactivity in either group. Alternatively, the higher dose could have caused an interaction with other neurotransmitter pathways. While PCA has a relatively high affinity for the 5-HT system, it also increases dopamine release at higher doses (Henderson *et al.* 1993). Therefore, PCA co-administered with DA receptor antagonists could provide valuable data to better understand these interactions. Additionally, use of a more specific serotonin agonist, such as 8-OH-DPAT, a compound that favors the 5-HT_1a_ receptors that are important for mood and depression-like behaviors (Cunha *et al.* 2013b), could provide a more nuanced finding.

The main findings of this study were that Cr deficiency leads to resiliency in LH and that a total loss of Cr causes augmentations to the serotonergic system without creating an endogenous depression- or anxiety-like phenotype. These mice show significant cognitive deficits (Skelton *et al.* 2011; Udobi *et al.* 2018), in agreement with humans with X-linked Crt deficiency (CTD) who have moderate to severe intellectual disability and delayed language development (van de Kamp *et al.* 2013a; van de Kamp *et al.* 2013b; van de Kamp *et al.* 2014; van de Kamp *et al.* 2012). While the cognitive phenotype of human CTD patients and *Slc6a8*^−/*y*^ mice are well-established, the results of this study provide a greater understanding of CTD by refining the behavioral profile of this important mouse model. Interestingly, humans with CTD are often described as having a “happy nature” and affective disorders have not been identified as a co-morbidity of CTD (van de Kamp *et al.* 2014). Therefore, results of this study further solidify this mouse as a high-fidelity model of CTD.

## Abbreviations

5-HT: serotonin
5-HIAA: 5-hydroxyindoleacetic acid
Cr: creatine
Crt: creatine transporter
EZM: elevated zero maze
HAM-D: Hamilton Depression Rating Scale
LH: learned helplessness
MAO: monoamine oxidase
PCA: *para*-chloroamphetamine
PCr: phosphocreatine
*Slc6a8*^−/*y*^: male *Slc6a8* knockout mice
*Slc6a8*^+/*y*^: male wild type mice
*Slc6a8*^+/−^: female mice heterozygous for *Slc6a8*
*Slc6a8*^+/+^: female wild type mice
SSRI: selective serotonin reuptake inhibitor
TPH-2: tryptophan hydroxylase-2
TST: tail-suspension test
cLH: rats bred for a congenital LH response

## Acknowledgements

The authors thank Marla K. Perna for editorial support. The authors declare no conflicts of interests.

## References

Allen, P. J. (2012) Creatine metabolism and psychiatric disorders: Does creatine supplementation have therapeutic value? Neurosci Biobehav Rev 36, 1442–1462.

Allen, P. J., D’Anci, K. E., Kanarek, R. B. and Renshaw, P. F. (2012) Sex-specific antidepressant effects of dietary creatine with and without sub-acute fluoxetine in rats. Pharmacol Biochem Behav 101, 588–601.

Amos-Kroohs, R. M., Williams, M. T., Braun, A. A., Graham, D. L., Webb, C. L., Birtles, T. S., Greene, R. M., Vorhees, C. V. and Pisano, M. M. (2013) Neurobehavioral phenotype of C57BL/6J mice prenatally and neonatally exposed to cigarette smoke. Neurotoxicol Teratol 35, 34–45.

Benjamini, Y., Krieger, A. M. and Yekutieli, D. (2006) Adaptive linear step-up procedures that control the false discovery rate. Biometrika 93, 491–507.

Braissant, O., Henry, H., Beard, E. and Uldry, J. (2011) Creatine deficiency syndromes and the importance of creatine synthesis in the brain. Amino Acids, 1315–1324.

Braissant, O., Henry, H., Loup, M., Eilers, B. and Bachmann, C. (2001) Endogenous synthesis and transport of creatine in the rat brain: an in situ hybridization study. Molecular Brain Research 86, 193–201.

Can, A., Dao, D. T., Chantelle E, T., Piantadosi, S. C., Bhat, S. and Gould, T. D. (2012) The Tail Suspension Test. Journal of Visualized Experiments.

Chourbaji, S., Zacher, C., Sanchis-Segura, C., Dormann, C., Vollmayr, B. and Gass, P. (2005) Learned helplessness: validity and reliability of depressive-like states in mice. Brain Res Brain Res Protoc 16, 70–78.

Coplan, J. D., Mathew, S. J., Mao, X., Smith, E. L., Hof, P. R., Coplan, P. M., Rosenblum, L. A., Gorman, J. M. and Shungu, D. C. (2006) Decreased choline and creatine concentrations in centrum semiovale in patients with generalized anxiety disorder: relationship to IQ and early trauma. Psychiatry Res 147, 27–39.

Cunha, M. P., Budni, J., Ludka, F. K. et al. (2016) Involvement of PI3K/Akt Signaling Pathway and Its Downstream Intracellular Targets in the Antidepressant-Like Effect of Creatine. Mol Neurobiol 53, 2954–2968.

Cunha, M. P., Budni, J., Pazini, F. L., Oliveira, A., Rosa, J. M., Lopes, M. W., Leal, R. B. and Rodrigues, A. L. (2014) Involvement of PKA, PKC, CAMK-II and MEK1/2 in the acute antidepressant-like effect of creatine in mice. Pharmacol Rep 66, 653–659.

Cunha, M. P., Machado, D. G., Capra, J. C., Jacinto, J., Bettio, L. E. and Rodrigues, A. L. (2012) Antidepressant-like effect of creatine in mice involves dopaminergic activation. J Psychopharmacol 26, 1489–1501.

Cunha, M. P., Pazini, F. L., Ludka, F. K. et al. (2015) The modulation of NMDA receptors and L-arginine/nitric oxide pathway is implicated in the anti-immobility effect of creatine in the tail suspension test. Amino Acids 47, 795–811.

Cunha, M. P., Pazini, F. L., Oliveira, A., Bettio, L. E., Rosa, J. M., Machado, D. G. and Rodrigues, A. L. (2013a) The activation of alpha1-adrenoceptors is implicated in the antidepressant-like effect of creatine in the tail suspension test. Prog Neuropsychopharmacol Biol Psychiatry 44, 39–50.

Cunha, M. P., Pazini, F. L., Oliveira, A., Machado, D. G. and Rodrigues, A. L. (2013b) Evidence for the involvement of 5-HT1A receptor in the acute antidepressant-like effect of creatine in mice. Brain Res Bull 95, 61–69.

Czeh, B., Michaelis, T., Watanabe, T., Frahm, J., Biurrun, G. d., Kampen, M. v., Bartolomucci, A. and Fuchs, E. (2001) Stress-induced changes in cerebral metabolites, hippocampal volume, and cell proliferation are prevented by antidepressant treatment with tianeptine. Proc Natl Acad Sci 98, 12796–12801.

Fuller, R. W. (1992) Effects of p-Chloroamphetamine on Brain Serotonin Neurons. Neurochemical Research 17, 449–456.

Gardner, A. and Boles, R. G. (2011) Beyond the serotonin hypothesis: mitochondria, inflammation and neurodegeneration in major depression and affective spectrum disorders. Prog Neuropsychopharmacol Biol Psychiatry 35, 730–743.

Geyer, M. A. (1996) Serotonergic functions in arousal and motor activity. Behavioural Brain Research, 31–35.

Hammels, C., Pishva, E., De Vry, J. et al. (2015) Defeat stress in rodents: From behavior to molecules. Neurosci Biobehav Rev 59, 111–140.

Hanna-El-Daher, L. and Braissant, O. (2016) Creatine synthesis and exchanges between brain cells: What can be learned from human creatine deficiencies and various experimental models? Amino Acids 48, 1877–1895.

Hautman, E. R., Kokenge, A. N., Udobi, K. C., Williams, M. T., Vorhees, C. V. and Skelton, M. R. (2014) Female mice heterozygous for creatine transporter deficiency show moderate cognitive deficits. J Inherit Metab Dis 37, 63–68.

Henderson, M. G., Perry, K. W. and Fuller, R. W. (1993) Possible Involvement of Dopamine in the Long Term Serotonin Depletion by p-Chloroamphetamine and beta,beta-Difluoro p-Chloroamphetamine in Rats. The Journal of Pharmacology and Experimental Therapeutics 267, 417–424.

Joncquel-Chevalier Curt, M., Voicu, P. M., Fontaine, M. et al. (2015) Creatine biosynthesis and transport in health and disease. Biochimie 119, 146–165.

Klinedinst, N. J. and Regenold, W. T. (2015) A mitochondrial bioenergetic basis of depression. J Bioenerg Biomembr 47, 155–171.

Lugenbiel, P., Sartorius, A., Vollmayr, B. and Schloss, P. (2010) Creatine transporter expression after antidepressant therapy in rats bred for learned helplessness. The World Journal of Biological Psychiatry 11, 329–333.

Lyoo, I. K., Yoon, S., Kim, T.-S., Hwang, J., Kim, J. E., Won, W., Bae, S. and Renshaw, P. F. (2012) A Randomized, Double-Blind Placebo-Controlled Trial of Oral Creatine Monohydrate Augmentation for Enhanced Response to a Selective Serotonin Reuptake Inhibitor in Women With Major Depressive Disorder. Am J Pyschiatry, 937–945.

Maier, S. F. and Seligman, M. E. (2016) Learned helplessness at fifty: Insights from neuroscience. Psychol Rev 123, 349–367.

NIMH (2010a) Top 10 Leading Disease/Disorder Categories Contributing to U.S. DALYs.

NIMH (2010b) U.S. DALYs for Mental and Behavioral Disorders as a Percent of Total U.S. DALYs.

Pazini, F. L., Cunha, M. P., Rosa, J. M., Colla, A. R., Lieberknecht, V., Oliveira, A. and Rodrigues, A. L. (2016) Creatine, Similar to Ketamine, Counteracts Depressive-Like Behavior Induced by Corticosterone via PI3K/Akt/mTOR Pathway. Mol Neurobiol 53, 6818–6834.

Perna, M. K., Kokenge, A. N., Miles, K. N., Udobi, K. C., Clark, J. F., Pyne-Geithman, G. J., Khuchua, Z. and Skelton, M. R. (2016) Creatine transporter deficiency leads to increased whole body and cellular metabolism. Amino Acids 48, 2057–2065.

Reed, A. L., Anderson, J. C., Bylund, D. B., Petty, F., El Refaey, H. and Happe, H. K. (2009) Treatment with escitalopram but not desipramine decreases escape latency times in a learned helplessness model using juvenile rats. Psychopharmacology (Berl) 205, 249–259.

Roitman, S., Green, T., Osher, Y., Karni, N. and Levine, J. (2007) Creatine monohydrate in resistant depression: a preliminary study. Bipolar Disorders, 754–758.

Sartorius, A., Vollmayr, B., Neumann-Haefelin, C., Ende, G., Hoehn, M. and Henn, F. A. (2003) Specific creatine rise in learned helplessness induced by electroconvulsive shock treatment. Neuroreport 14, 2199–2201.

Schaefer, T. L., Vorhees, C. V. and Williams, M. T. (2009) Mouse plasmacytoma-expressed transcript 1 knock out induced 5-HT disruption results in a lack of cognitive deficits and an anxiety phenotype complicated by hypoactivity and defensiveness. Neuroscience 164, 1431–1443.

Schuff, N., Neylan, T. C., Fox-Bosetti, S., Lenoci, M., Samuelson, K. W., Studholme, C., Kornak, J., Marmar, C. R. and Weiner, M. W. (2008) Abnormal N-acetylaspartate in hippocampus and anterior cingulate in posttraumatic stress disorder. Psychiatry Res 162, 147–157.

Skelton, M. R., Schaefer, T. L., Graham, D. L., Degrauw, T. J., Clark, J. F., Williams, M. T. and Vorhees, C. V. (2011) Creatine transporter (CrT; Slc6a8) knockout mice as a model of human CrT deficiency. PLoS One 6, e16187.

Sprowles, J. L., Hufgard, J. R., Gutierrez, A., Bailey, R. A., Jablonski, S. A., Williams, M. T. and Vorhees, C. V. (2016) Perinatal exposure to the selective serotonin reuptake inhibitor citalopram alters spatial learning and memory, anxiety, depression, and startle in Sprague-Dawley rats. Int J Dev Neurosci 54, 39–52.

Stockebrand, M., Nejad, A. S., Neu, A., Kharbanda, K. K., Sauter, K., Schillemeit, S., Isbrandt, D. and Choe, C. U. (2016) Transcriptomic and metabolic analyses reveal salvage pathways in creatine-deficient AGAT(-/-) mice. Amino Acids 48, 2025–2039.

Udobi, K. C., Kokenge, A. N., Hautman, E. R., Ullio, G., Coene, J., Williams, M. T., Vorhees, C. V., Mabondzo, A. and Skelton, M. R. (2018) Cognitive deficits and increases in creatine precursors in a brain-specific knockout of the creatine transporter gene Slc6a8. Genes Brain Behav 17, e12461.

van de Kamp, J. M., Betsalel, O. T., Mercimek-Mahmutoglu, S. et al. (2013a) Phenotype and genotype in 101 males with X-linked creatine transporter deficiency. J Med Genet 50, 463–472.

van de Kamp, J. M., Jakobs, C., Gibson, K. M. and Salomons, G. S. (2013b) New insights into creatine transporter deficiency: the importance of recycling creatine in the brain. J Inherit Metab Dis 36, 155–156.

van de Kamp, J. M., Mancini, G. M. and Salomons, G. S. (2014) X-linked creatine transporter deficiency: clinical aspects and pathophysiology. J Inherit Metab Dis 37, 715–733.

van de Kamp, J. M., Pouwels, P. J., Aarsen, F. K. et al. (2012) Long-term follow-up and treatment in nine boys with X-linked creatine transporter defect. J Inherit Metab Dis 35, 141–149.

van der Hart, M. G., Czeh, B., de Biurrun, G., Michaelis, T., Watanabe, T., Natt, O., Frahm, J. and Fuchs, E. (2002) Substance P receptor antagonist and clomipramine prevent stress-induced alterations in cerebral metabolites, cytogenesis in the dentate gyrus and hippocampal volume. Mol Psychiatry 7, 933–941.

Villarreal, G., Petropoulos, H., Hamilton, D. A., Rowland, L. M., Horan, W. P., Griego, J. A., Moreshead, M., Hart, B. L. and Brooks, W. M. (2002) Proton Magnetic Resonance Spectroscopy of the Hippocampus and Occipital White Matter in PTSD: Preliminary Results. Can J Psychiatry 47, 666–670.

Wyss, M., Boles, R. G. (2000) Creatine and Creatinine Metabolism. Physiol Rev 3, 1107–1213.

